# Enhancing the Clinical Utility of Renal Denervation Through Efficient Physiologic Monitoring

**DOI:** 10.64898/2026.09.23.753680

**Authors:** Kari Shad, Alicia M. Schiller, Irving H. Zucker, Yiannis S. Chatzizisis, Han-Jun Wang, Peter Ricci Pellegrino

## Abstract

**Introduction:** The inability to measure renal sympathetic outflow clinically poses a significant problem for patient identification and intraprocedural feedback for renal denervation. Sympathetic vasomotion is a novel physiologic measure of rhythmic vascular control that reflects renal sympathetic nervous system activity, but previous measurements of sympathetic vasomotion relied on 15-minute data recordings, which may be impractical in many clinical settings. We therefore tested whether shorter data segments can provide reliable sympathetic vasomotion assessment to enhance decision making for renal denervation.

**Methods:** Ten swine that had previously undergone unilateral surgical renal denervation underwent four rounds of catheter-based denervation of the contralateral kidney. Sympathetic vasomotion was quantified from the time-varying arterial pressure-resistive renal blood flow transfer function of recordings obtained before and after each round of ablation. Randomly placed 1- to 14-minute data segments were scored and compared with the corresponding 15-minute sympathetic vasomotion scores across 100 passes using correlation, intraclass correlation, error, and Bland-Altman agreement metrics, and by whether they reproduced the denervation decisions made from the full recordings.

**Results:** Agreement with the 15-minute reference score improved steeply with recording length up to approximately four minutes and slowly thereafter, with the point of diminishing returns at four minutes for every group-level measure. Four-minute data segments showed an intraclass correlation coefficient of 0.94, reproduced the group-level statistical results across ablations, and preserved intraprocedural denervation decisions. Bland-Altman analysis revealed that individual-level precision continued to improve with recording length; the proportion of scores falling within 25 units of their own 15-minute score was 65 percent at four minutes and 96 percent at ten minutes.

**Discussion:** Four-minute data segments provide a feasible minimum recording duration when the question is a group comparison or a paired, within-individual decision, such as intraprocedural confirmation that an innervated kidney has shifted toward a denervated state. When a single score must be interpreted on its own, as in preprocedural patient selection, ten-minute recordings provide greater precision. Further studies are needed to prospectively evaluate sympathetic vasomotion in clinical settings and determine whether this technology can support procedural feedback and inform patient selection.

## Introduction

Hypertension affects nearly half of adults in the United States and is the leading modifiable risk factor for cardiovascular disease (Mensah et al., 2023). Multisociety guidelines strongly recommend targeting systolic blood pressure less than 130 mm Hg and diastolic blood pressure less than 80 mm Hg in patients at risk for cardiovascular disease (Jones et al., 2025). For decades, pharmacotherapy has been the therapeutic mainstay for hypertension, but problems with efficacy, tolerance, and adherence have led to renewed interest in interventional treatments for hypertension (Pimenta and Calhoun, 2012; Şener et al., 2026).

The leading interventional antihypertensive therapy is renal denervation, an intravascular catheter-based ablation procedure targeting the renal sympathetic nerves (DiBona and Esler, 2010). The initial renal denervation clinical trials showed dramatic blood pressure reductions in patients with treatment-resistant hypertension, creating a groundswell of enthusiasm and spurring industry investment (Krum et al., 2009; Esler et al., 2010). This momentum was stymied after the first sham-controlled trial failed to meet its primary efficacy endpoint (Bhatt et al., 2014). Post-hoc analyses implicated inadequate renal denervation and the lack of intraprocedural feedback as key factors in this failure (Esler, 2014; Kandzari et al., 2015). More recent sham-controlled trials with second-generation devices designed to more comprehensively ablate the renal sympathetic nerves have generally shown positive results, leading to the United States Food and Drug Administration approval of two devices in 2023 (Azizi et al., 2018, 2022; Böhm et al., 2020). Despite these advances in device design, procedural success remains difficult to confirm in real time because there is no reliable intraprocedural physiologic marker to confirm adequate denervation (Paton et al., 2026). Accurately measuring renal sympathetic nervous system activity during these procedures could improve confirmation of procedural success and strengthen the reliability of future studies evaluating the efficacy of renal denervation for the treatment of hypertension (Gulati et al., 2016).

Similarly, the lack of markers of renal sympathetic outflow complicates patient selection for this invasive procedure (Paton et al., 2026). Hypertension is a heterogeneous disorder, and hypertensive patients exhibit variability in regional sympathetic outflow when assessed by norepinephrine spillover (Esler et al., 1989). Parameters associated with increased sympathetic activity, including heart rate and plasma renin activity, have been associated with renal denervation responsiveness in some cohorts, underscoring the potential that variability in renal sympathetic tone underlies the variable antihypertensive effect of renal denervation (Schmieder et al., 2024). A more specific preprocedural measure of renal sympathetic outflow could thereby help identify patients most likely to benefit from this procedure and maximize the clinical value of this technique.

To address the lack of a clinically implementable marker for renal sympathetic outflow, our group identified a novel measure, sympathetic vasomotion, which refers to rhythmic fluctuations in sympathetic vascular control that can be detected through simultaneous measurements of arterial pressure and blood flow. The ubiquity of clinical technology to measure arterial pressure and blood flow makes this a practical physiologic marker of renal sympathetic tone and procedural response during renal denervation (Pellegrino et al., 2020, 2025).

Previous work from our group, however, relied on quantification of 15 minutes of continuous arterial pressure and blood flow data. Although feasible in a preclinical research setting, 15-minute recording periods are impractical in the cardiac catheterization laboratory, where procedural time carries substantial clinical and financial implications. Moreover, obtaining 15 minutes of artifact-free continuous renal blood flow velocity with transabdominal ultrasound, as might be done to identify patients preprocedurally with high renal sympathetic tone, poses a technical challenge for even highly skilled ultrasonographers. Thus, this constraint limits the clinical application of sympathetic vasomotion as an intraprocedural and preprocedural monitoring tool. To address this limitation, we evaluated the reliability of shorter data segments for measuring sympathetic vasomotion, with the goal of identifying a duration that preserves measurement accuracy while minimizing acquisition time for different clinical contexts.

## Materials and Methods

### Data and Materials Availability

Data and source code used for this study are available on figshare (doi: 10.6084/m9.figshare.33274521). The corresponding author is responsible for the maintenance of this repository.

### Swine Experiments

This study analyzed hemodynamic data that were described in detail in a previous publication (Pellegrino et al., 2025). In brief, 10 swine first underwent unilateral surgical renal denervation (SDNx), providing a denervated comparison (Figure 1A). One week later, each animal subsequently underwent catheter-based renal denervation (CDNx) of the contralateral kidney using an FDA-approved radiofrequency ablation system in a regimented distal-to-proximal fashion: first the largest renal branch artery was ablated, followed by the remaining branch arteries, followed by the distal main renal artery, and finally the proximal main renal artery. Bilateral renal blood flow velocity and arterial pressure were recorded for 15 minutes before and after these four rounds of radiofrequency ablation targeting the renal sympathetic nerves. All procedures were reviewed and approved by the University of Nebraska Medical Center Institutional Animal Care and Use Committee and carried out in accordance with the NIH Guide for the Care and Use of Laboratory Animals.

**Fig. 1.**
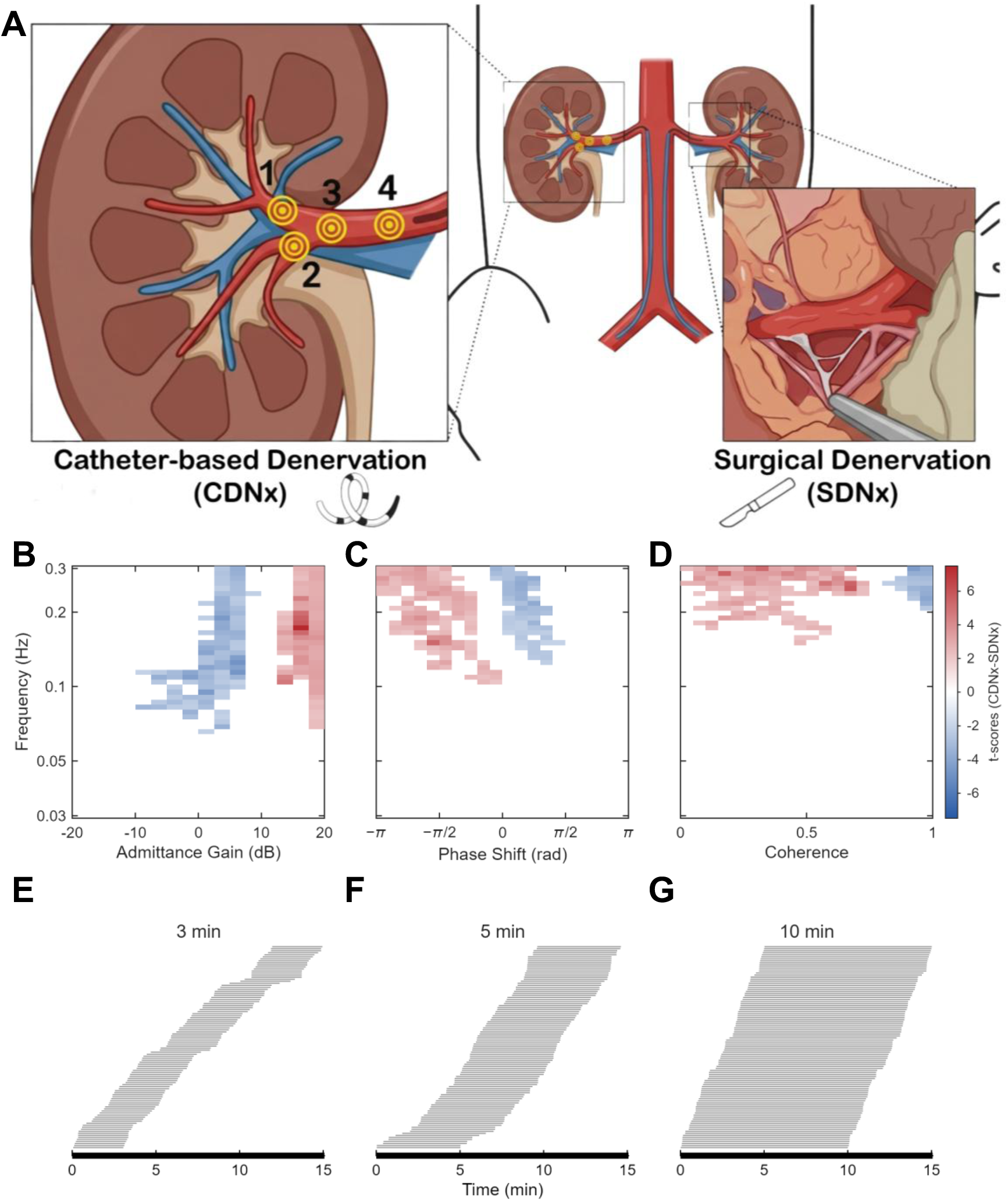
Experimental paradigm and quantification of sympathetic vasomotion. (A) Surgical renal denervation (SDNx) was performed on one kidney and four rounds of catheter-based renal denervation (CDNx) were performed on the contralateral kidney, with measurement of arterial pressure and renal blood flow velocity before and after each round of ablation. (B-D) Sympathetic vasomotion cluster maps for (B) admittance gain, (C) phase shift, and (D) coherence of the time-varying pressure-resistive flow transfer function at baseline (n = 10 swine). Each cell is one frequency (y-axis, logarithmic) and one value bin (x-axis) of the percent-occurrence distribution. Color, on one scale shared by B through D, shows the paired t-score comparing occupancy in the CDNx kidney, innervated at baseline, with the SDNx kidney: red cells were occupied more often by the CDNx kidney and blue cells by the SDNx kidney. Cells with an absolute t-score greater than 2 were grouped into connected clusters, and the largest positive and largest negative cluster in each channel were retained (166, 169, and 136 cells for admittance gain, phase shift, and coherence). Other cells are blank. A kidney’s sympathetic vasomotion score is the sum, over retained cells, of the t-score multiplied by that kidney’s occupancy, scaled so that the mean baseline CDNx kidney scores 100 and the mean baseline SDNx kidney scores 0. (E-G) Segment analysis. For each 15-minute recording (thick horizontal axis), 100 data segments of (E) 3, (F) 5, or (G) 10 minutes were placed at random start times (thin gray lines, sorted by start time) and each was scored and compared with the score from the full recording. The data segments shown are those used for one animal’s baseline recording.

### Sympathetic Vasomotion Scoring

The arterial pressure and bilateral renal blood flow velocity data were then used to calculate sympathetic vasomotion scores from the t-statistic difference cluster maps derived from the time-varying arterial pressure-resistive renal blood flow transfer function occurrence as described previously (Pellegrino et al., 2025). In brief, the score is a weighted sum over three channels of the product of the occurrence and the t-score weight, with 0 representing the average sympathetic vasomotion in a denervated kidney and 100 representing the average sympathetic vasomotion in an innervated kidney (Figure 1B-D).

### Data Analysis

Analysis of hemodynamic data was performed in MATLAB 2025a (MathWorks, Inc., Natick, Massachusetts, USA). The 15-minute sympathetic vasomotion score was treated as the reference value for each animal, condition, and ablation number. To determine whether shorter data recordings could reproduce the 15-minute sympathetic vasomotion score, vasomotion scores from shortened data segments ranging from 1 to 14 minutes were compared with the corresponding 15-minute data segment. For each shortened data segment duration, 100 randomized passes were performed (Figure 1E-G). Analyses were performed across catheter denervated and surgically denervated kidneys and across ablations.

### Statistical Analysis

Group-level agreement was assessed to determine whether shortened data segments preserved the overall relationships observed with the 15-minute sympathetic vasomotion scores. For each data segment duration and randomized pass, shortened data segment sympathetic vasomotion scores were compared with the corresponding 15-minute sympathetic vasomotion scores using intraclass correlation coefficient (ICC), Pearson correlation, Spearman correlation, root mean square error (RMSE), and mean absolute error (MAE). Metrics were summarized across the 100 randomized passes for each data segment duration using the median and 2.5th to 97.5th percentile range. The optimal duration for group-level agreement was assessed both by the absolute values of these parameters and by the point of diminishing returns in these parameters with increasing data segment length using the Kneedle method (Satopää et al., 2011; Koo and Li, 2016).

To assess the ability to make statistical inferences with shorter data segments, repeated-measures analysis of variance (RM-ANOVA) with denervation modality and ablation round as within-subjects factors with Greenhouse-Geisser correction for sphericity was performed on the 100 passes of shorter data segments to test the consistency with which this reproduced the observed statistical results from the 15-minute data. The median P values for the RM-ANOVA terms and the percentage of passes for which these P values reached statistical significance were calculated.

Individual-level agreement was assessed to determine whether a shortened data segment could be used interchangeably with the 15-minute sympathetic vasomotion score for individual observations. For each segment duration and randomized pass, Bland-Altman metrics were calculated from the difference between the shortened data segment sympathetic vasomotion score and the corresponding 15-minute sympathetic vasomotion score. Bias was calculated as the mean difference, absolute bias as the absolute value of the mean difference, and the standard deviation (SD) of the differences was used to calculate the limits of agreement (LOA) as bias +/- 1.96 x SD. The percentage of observations falling within 25 sympathetic vasomotion score units (i.e., one quarter of the difference between innervated and denervated kidneys at baseline) of their 15-minute reference score was also calculated for each segment duration. Additionally, representative Bland-Altman plots were generated for one representative (SD equal to the median) pass of three durations (3 minutes, 5 minutes, and 10 minutes) to visualize individual-level agreement with the 15-minute sympathetic vasomotion score.

The effect of recording segment length on decision-making performance was assessed through three parameters. First, the percentage of correct kidney classifications prior to catheter-based renal denervation (i.e., sympathetic vasomotion scores showing CDNx greater than SDNx at baseline) was computed for various data segment lengths.

Second, the area under the curve for kidney classification as a function of sympathetic vasomotion score was calculated for recordings from 1-14 minutes. Finally, the area under the curve for the change in sympathetic vasomotion after both two (complete branch denervation) and four (complete) rounds of catheter-based radiofrequency ablation was calculated as a function of data segment duration.

## Results

### Group-Level Agreement

Agreement between shortened data segments and the 15-minute reference score was assessed across 100 randomized passes at each duration from 1 to 14 minutes (Table 1, Figure 2A and 2B). Agreement remained high down to four minutes (ICC 0.94, Pearson 0.95, RMSE 28.4, MAE 22.3 score units). Below four minutes, agreement declined more rapidly to the point that one-minute data segments had an ICC of 0.73 with an RMSE of 63.7 score units. These results indicate that data segments of approximately four minutes preserved group-level relationships, whereas further shortening resulted in a pronounced loss of agreement.

**Fig. 2.**
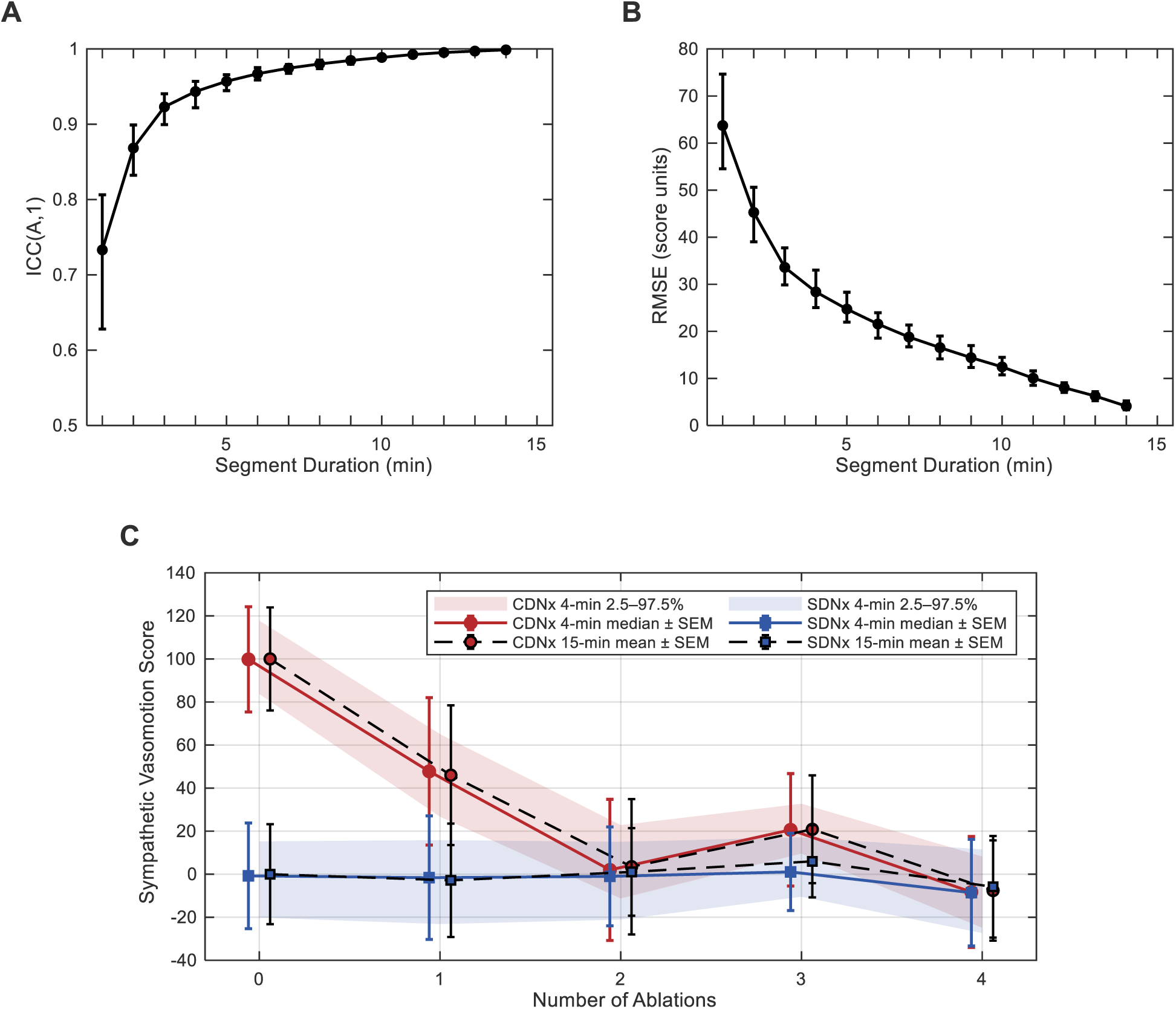
Group-level agreement between shortened data segments and the 15-minute sympathetic vasomotion score. (A) Intraclass correlation coefficient (ICC[A,1]; two-way, absolute agreement, single measures) and (B) root mean square error (RMSE) between data segment and 15-minute scores, for data segment durations of 1 to 14 minutes. For each duration, one data segment was placed at random in every recording and compared with that recording’s 15-minute score, pooling all 100 kidney-timepoints (10 animals x 5 timepoints x 2 kidneys). This was repeated 100 times. Lines show the median across the 100 repetitions and error bars the 2.5th to 97.5th percentiles. Agreement rose steeply up to about 4 minutes (ICC 0.94, RMSE 28.4 score units), the knee of the curve (Table 2), and more slowly thereafter (10 minutes: ICC 0.99, RMSE 12.5). (C) Sympathetic vasomotion scores of the catheter-based denervation (CDNx) kidney (red, circles), innervated at baseline, and the surgically denervated (SDNx) kidney (blue, squares), at baseline (0) and after each of four rounds of radiofrequency ablation (n = 10 swine). Black dashed lines show the 15-minute mean plus or minus SEM. Colored solid lines show the 4-minute scores as the median, across 100 sets of randomly placed 4-minute data segments, of the group mean plus or minus SEM. Shaded bands span the 2.5th to 97.5th percentiles of the 4-minute group mean across those 100 sets.

**Table 1:** Group-level agreement between shortened data segments and the 15-minute data segments across 100 randomized passes. In each pass, one data segment was placed at random in every recording and compared with that recording’s 15-minute score, pooling all 100 kidney-timepoints (10 swine x 5 timepoints x 2 kidneys). All values are the median over the 100 passes (2.5th to 97.5th percentile). As data segment duration decreased, agreement became less consistent, with lower intraclass correlation coefficient (ICC[A,1]), Pearson correlation, and Spearman correlation values and higher root mean square error (RMSE) and mean absolute error (MAE) at shorter durations. Overall, these results suggest that shortened data segments preserve strong group-level agreement, whereas the shortest durations show progressively greater disagreement from the 15-minute reference data.

| Duration (min) | ICC(A,1) | Pearson r | Spearman r | RMSE | MAE |
| --- | --- | --- | --- | --- | --- |
| 1 | 0.73 (0.63-0.81) | 0.74 (0.63-0.82) | 0.72 (0.63-0.81) | 63.7 (54.5-74.7) | 50.3 (43.6-59.8) |
| 2 | 0.87 (0.83-0.90) | 0.88 (0.84-0.91) | 0.86 (0.82-0.90) | 45.3 (39.0-50.6) | 35.7 (30.8-40.1) |
| 3 | 0.92 (0.90-0.94) | 0.93 (0.90-0.95) | 0.92 (0.89-0.94) | 33.6 (29.9-37.7) | 26.7 (23.5-31.2) |
| 4 | 0.94 (0.92-0.96) | 0.95 (0.93-0.96) | 0.94 (0.91-0.95) | 28.4 (25.1-33.0) | 22.3 (19.1-25.2) |
| 5 | 0.96 (0.94-0.97) | 0.96 (0.95-0.97) | 0.95 (0.93-0.96) | 24.7 (22.0-28.3) | 18.9 (16.6-21.5) |
| 6 | 0.97 (0.96-0.98) | 0.97 (0.96-0.98) | 0.96 (0.95-0.97) | 21.6 (18.6-24.0) | 16.5 (14.5-18.5) |
| 7 | 0.97 (0.97-0.98) | 0.98 (0.97-0.98) | 0.97 (0.96-0.98) | 18.8 (16.7-21.4) | 14.3 (12.6-16.3) |
| 8 | 0.98 (0.97-0.99) | 0.98 (0.97-0.99) | 0.97 (0.96-0.98) | 16.5 (14.2-19.0) | 12.7 (11.2-14.3) |
| 9 | 0.98 (0.98-0.99) | 0.99 (0.98-0.99) | 0.98 (0.97-0.99) | 14.4 (12.3-17.0) | 11.1 (9.7-12.8) |
| 10 | 0.99 (0.98-0.99) | 0.99 (0.99-0.99) | 0.98 (0.98-0.99) | 12.5 (10.8-14.5) | 9.6 (8.0-10.6) |
| 11 | 0.99 (0.99-0.99) | 0.99 (0.99-0.99) | 0.99 (0.98-0.99) | 10.1 (8.5-11.6) | 7.8 (6.7-9.0) |
| 12 | 1.00 (0.99-1.00) | 1.00 (0.99-1.00) | 0.99 (0.99-0.99) | 8.0 (7.0-9.1) | 6.2 (5.5-7.0) |
| 13 | 1.00 (1.00-1.00) | 1.00 (1.00-1.00) | 1.00 (0.99-1.00) | 6.3 (5.3-7.2) | 4.8 (4.1-5.4) |
| 14 | 1.00 (1.00-1.00) | 1.00 (1.00-1.00) | 1.00 (1.00-1.00) | 4.1 (3.3-5.2) | 3.0 (2.6-3.4) |

**Table 2:** Point of diminishing returns for each group-level agreement measure. For each measure and each of the 100 passes, the knee of the measure-duration curve (1 to 14 minutes) was located with the Kneedle method: both axes were rescaled to 0 to 1, and the knee was taken as the duration at which the rescaled curve lies furthest above the chord joining its endpoints. RMSE and MAE, which fall with duration, were negated first. The knee is shown as the median across passes (2.5th to 97.5th percentile), together with the most common knee and the percentage of passes at that value, and the measure at each pass’s own knee. All five measures placed the point of diminishing returns at approximately four minutes.

| Measure | Knee (min) | Modal knee (min, % of passes) | Value at knee |
| --- | --- | --- | --- |
| ICC(A,1) | 4 (2-5) | 4 (45%) | 0.94 (0.91-0.96) |
| Pearson r | 3.5 (2-5) | 3 (45%) | 0.94 (0.90-0.96) |
| Spearman r | 4 (2-5) | 4 (42%) | 0.94 (0.91-0.96) |
| RMSE | 4 (3-5) | 4 (52%) | 27.3 (21.1-34.9) |
| MAE | 4 (3-6) | 4 (49%) | 21.0 (15.5-25.8) |

### Point of Diminishing Returns

Because agreement continued to improve with recording length, the point of diminishing returns was identified from the knee of each measure-duration curve using the Kneedle method, computed separately for each of the 100 passes (Table 2). This analysis identified knees at 4 minutes for ICC(A,1), Spearman correlation, RMSE, and MAE, and at 3.5 minutes for the Pearson correlation. For all of the 100 passes, these knees fell between 2 and 6 minutes. Lengthening the recording beyond this point continued to improve agreement, but each additional minute added less than 0.015 to the ICC.

### Group-Level Statistical Inferences

Four-minute data segments also preserved the overall pattern observed with the 15-minute sympathetic vasomotion scores across ablations (Figure 2C, Table 3). Median four-minute catheter-based denervation (CDNx) scores closely tracked the 15-minute CDNx means, at 100, 48, 2, 21, and -8 score units at baseline and after each of the four ablations, against 100, 46, 3, 21, and -8 score units with the full recordings. Both showed a marked decrease in sympathetic vasomotion score from baseline through the first two ablations, followed by little additional reduction with subsequent ablations. The surgical denervation (SDNx) median likewise followed the 15-minute SDNx mean, remaining near zero during contralateral denervation. In the two-way RM-ANOVA, the interaction between denervation modality and ablation round was significant in 84 percent of 4-minute passes and 100 percent of 8-minute passes, indicative of adequate statistical power for cohort analysis at abbreviated lengths. Moreover, ablation round was not statistically significant for any of the passes for data segments 2 minutes or longer, indicative of group-level stability over time and protection against type I statistical errors. This indicates that shortened data segments are sensitive enough to detect true group-level effects and reliable enough to avoid false positives.

**Table 3:** Group-level statistical inference as a function of recording length. A two-way repeated-measures ANOVA of denervation modality (CDNx, SDNx) by ablation round (0 to 4), with Greenhouse-Geisser correction, was run on each of 100 randomized passes at every data segment duration (n = 10 swine). Columns give the median p value across passes for the modality by ablation round interaction, the primary effect, and for each main effect, followed by the percentage of passes reaching p < 0.05 for the same three effects. The final row is the full 15-minute recording, which yields a single p value per effect. The interaction was recovered in most passes from three minutes onward and in every pass from eight minutes, the borderline main effect of modality required longer data segments, and the main effect of ablation round, absent at 15 minutes, was absent at every duration.

| Duration (min) | Median P, denervation modality x ablation round | Median P, denervation modality | Median P, ablation round | % Passes P < 0.05, denervation modality x ablation round | % Passes P < 0.05, denervation modality | % Passes P < 0.05, ablation round |
| --- | --- | --- | --- | --- | --- | --- |
| 1 | 0.089 | 0.099 | 0.393 | 31% | 27% | 3% |
| 2 | 0.029 | 0.059 | 0.336 | 61% | 45% | 0% |
| 3 | 0.022 | 0.049 | 0.316 | 69% | 51% | 0% |
| 4 | 0.018 | 0.049 | 0.307 | 84% | 51% | 0% |
| 5 | 0.016 | 0.039 | 0.279 | 84% | 64% | 0% |
| 6 | 0.013 | 0.040 | 0.309 | 97% | 76% | 0% |
| 7 | 0.011 | 0.038 | 0.307 | 97% | 75% | 0% |
| 8 | 0.009 | 0.038 | 0.320 | 100% | 80% | 0% |
| 9 | 0.011 | 0.037 | 0.301 | 100% | 88% | 0% |
| 10 | 0.010 | 0.037 | 0.299 | 100% | 94% | 0% |
| 11 | 0.010 | 0.039 | 0.283 | 100% | 92% | 0% |
| 12 | 0.009 | 0.040 | 0.289 | 100% | 94% | 0% |
| 13 | 0.010 | 0.042 | 0.283 | 100% | 97% | 0% |
| 14 | 0.010 | 0.047 | 0.280 | 100% | 78% | 0% |
| 15 | 0.010 | 0.055 | 0.285 | - | - | - |

### Individual-Level Agreement

Bland-Altman analysis demonstrated that the systematic bias between shortened data segments and the 15-minute reference data remained small at all data segment durations, whereas the variability of individual differences increased as data segment duration decreased (Table 4, Figure 3). The median bias was 6.8 sympathetic vasomotion score units at one minute and within one score unit of zero from two minutes onward. Median absolute bias decreased from 7.3 score units at one minute to 2.2 score units at four minutes and 1.0 score unit at ten minutes. Decreasing data segment duration therefore had little effect on systematic bias except at the shortest durations.

**Fig. 3.**
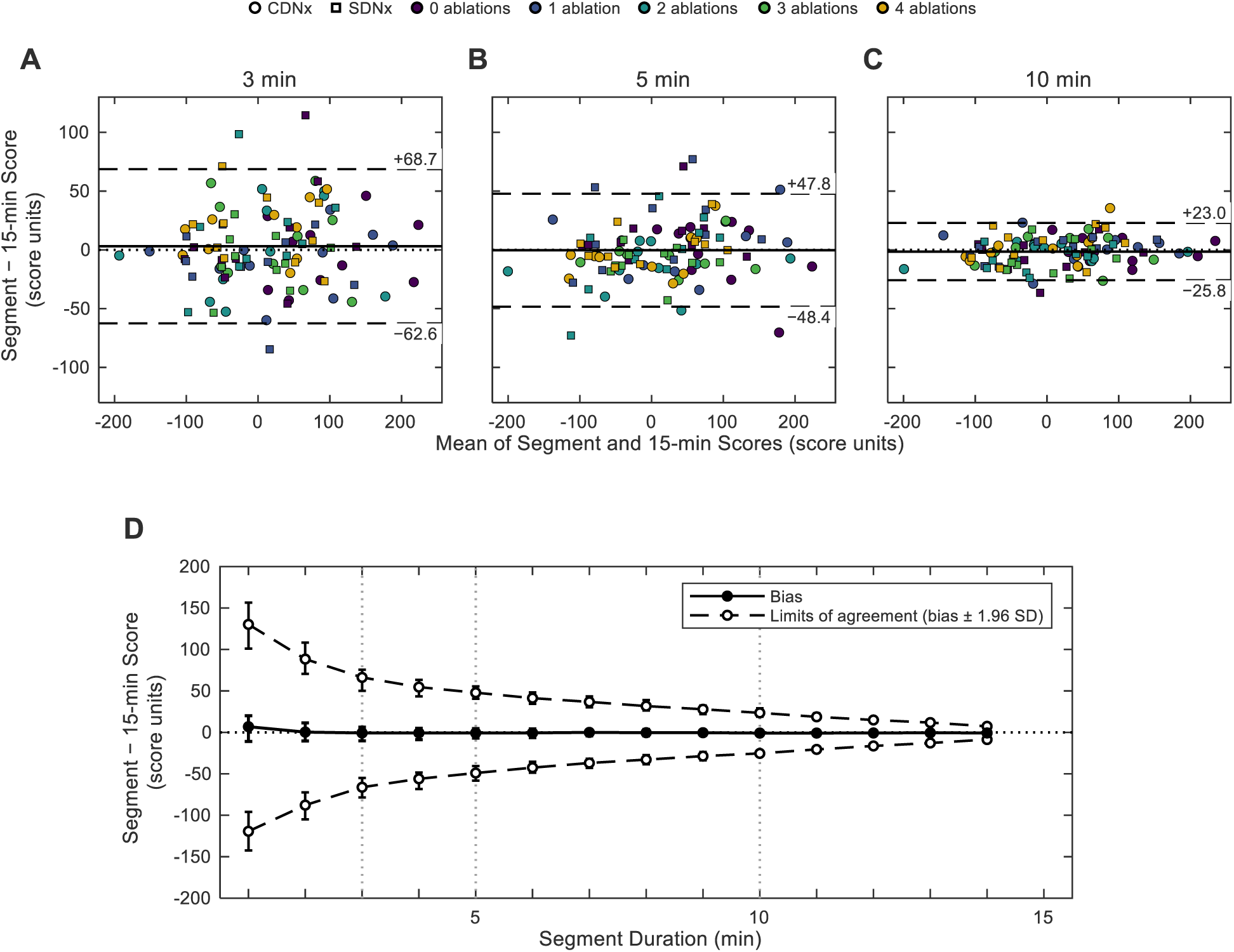
Individual-level agreement between shortened data segments and the 15-minute sympathetic vasomotion score. (A-C) Bland-Altman plots comparing (A) 3-minute, (B) 5-minute, and (C) 10-minute data segment scores with the corresponding 15-minute scores. Each point is one kidney at one timepoint (n = 100: 10 swine x 5 timepoints x 2 kidneys). Circles show the catheter-based denervation (CDNx) kidney and squares the surgically denervated (SDNx) kidney, and color indicates the number of ablations. The solid line marks the bias (mean difference) and dashed lines the limits of agreement (bias plus or minus 1.96 SD of the differences), with values at the right. Each plot shows a representative set of data segments: of 100 sets of randomly placed data segments at that duration, the one whose limits of agreement had the median width. The biases were +3.1 (3 min), -0.3 (5 min), and -1.4 (10 min) score units. (D) Bias (solid) and limits of agreement (dashed) against data segment duration from 1 to 14 minutes. Lines show the median across the 100 repetitions and error bars the 2.5th to 97.5th percentiles. Dotted vertical lines mark the durations shown in A through C. Bias was 6.8 score units at 1 minute and within 1 score unit of zero from 2 minutes onward. The limits of agreement narrowed from -119 to +130 score units at 1 minute to -56 to +55 at 4 minutes and -25 to +24 at 10 minutes.

**Table 4:**
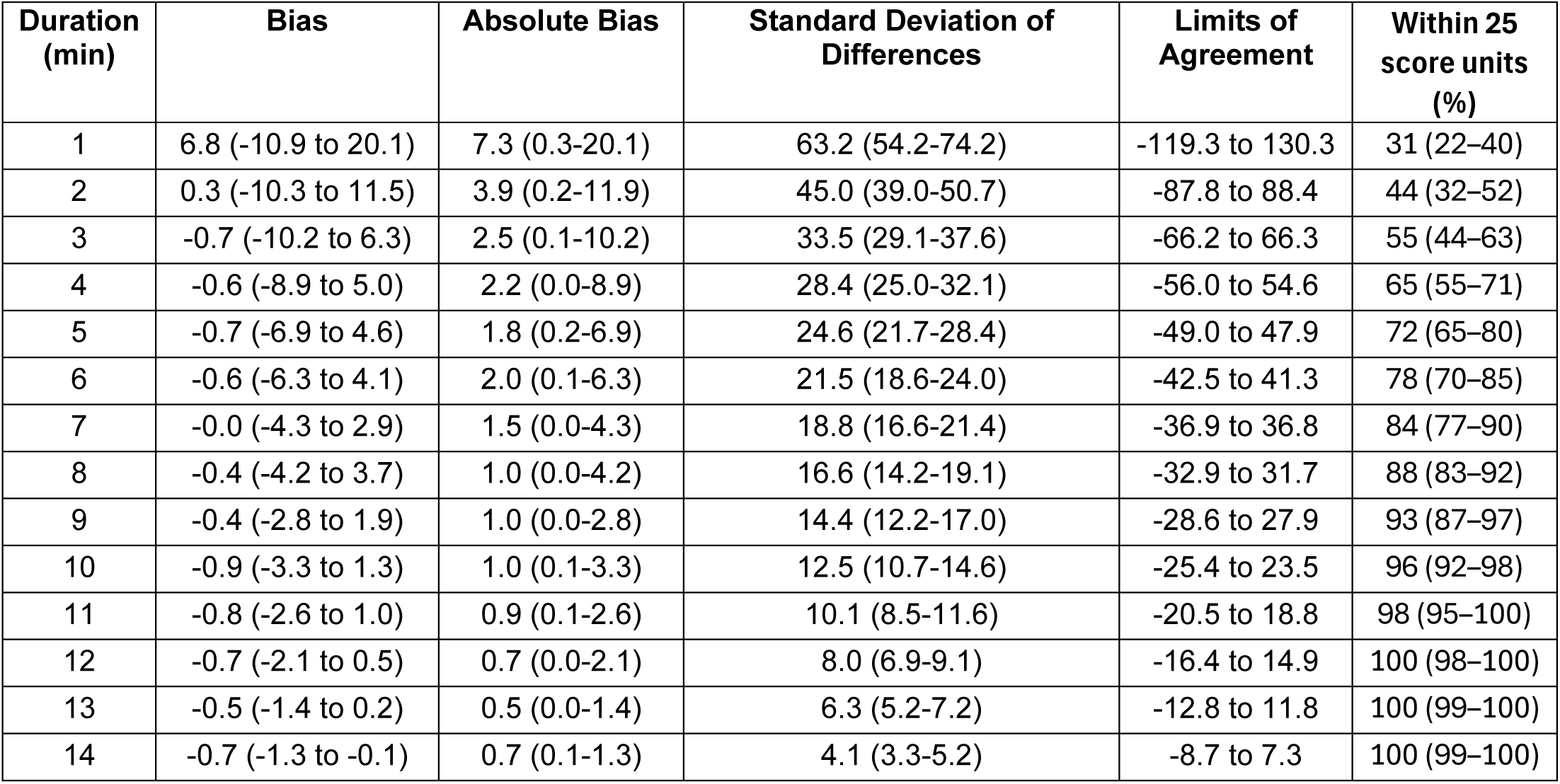
Bland-Altman agreement metrics between shortened data segments and 15-minute data segments across 100 randomized passes. Bias is the mean difference between the shortened data segment and the 15-minute score within a pass; absolute bias is the magnitude of that bias, and the limits of agreement are the bias plus or minus 1.96 times the standard deviation (SD) of the differences; within 25 score units represents the percentage of observations whose segment score fell within 25 units of their own 15-minute score. All values are the median over 100 passes (2.5th to 97.5th percentile). Median bias and absolute bias remained small across data segment durations, indicating limited systematic difference between the shortened and the 15-minute reference data. In contrast, the SD of the differences and the limits of agreement widened substantially as data segment duration decreased, reflecting greater variability in individual-level agreement with shorter data recordings. These findings indicate that reduced data segment duration has a modest effect on systematic bias but a larger effect on the precision of individual-level agreement.

| Duration (min) | Bias | Absolute Bias | Standard Deviation of Differences | Limits of Agreement | Within 25 score units (%) |
| --- | --- | --- | --- | --- | --- |
| 1 | 6.8 (-10.9 to 20.1) | 7.3 (0.3-20.1) | 63.2 (54.2-74.2) | -119.3 to 130.3 | 31 (22–40) |
| 2 | 0.3 (-10.3 to 11.5) | 3.9 (0.2-11.9) | 45.0 (39.0-50.7) | -87.8 to 88.4 | 44 (32–52) |
| 3 | -0.7 (-10.2 to 6.3) | 2.5 (0.1-10.2) | 33.5 (29.1-37.6) | -66.2 to 66.3 | 55 (44–63) |
| 4 | -0.6 (-8.9 to 5.0) | 2.2 (0.0-8.9) | 28.4 (25.0-32.1) | -56.0 to 54.6 | 65 (55–71) |
| 5 | -0.7 (-6.9 to 4.6) | 1.8 (0.2-6.9) | 24.6 (21.7-28.4) | -49.0 to 47.9 | 72 (65–80) |
| 6 | -0.6 (-6.3 to 4.1) | 2.0 (0.1-6.3) | 21.5 (18.6-24.0) | -42.5 to 41.3 | 78 (70–85) |
| 7 | -0.0 (-4.3 to 2.9) | 1.5 (0.0-4.3) | 18.8 (16.6-21.4) | -36.9 to 36.8 | 84 (77–90) |
| 8 | -0.4 (-4.2 to 3.7) | 1.0 (0.0-4.2) | 16.6 (14.2-19.1) | -32.9 to 31.7 | 88 (83–92) |
| 9 | -0.4 (-2.8 to 1.9) | 1.0 (0.0-2.8) | 14.4 (12.2-17.0) | -28.6 to 27.9 | 93 (87–97) |
| 10 | -0.9 (-3.3 to 1.3) | 1.0 (0.1-3.3) | 12.5 (10.7-14.6) | -25.4 to 23.5 | 96 (92–98) |
| 11 | -0.8 (-2.6 to 1.0) | 0.9 (0.1-2.6) | 10.1 (8.5-11.6) | -20.5 to 18.8 | 98 (95–100) |
| 12 | -0.7 (-2.1 to 0.5) | 0.7 (0.0-2.1) | 8.0 (6.9-9.1) | -16.4 to 14.9 | 100 (98–100) |
| 13 | -0.5 (-1.4 to 0.2) | 0.5 (0.0-1.4) | 6.3 (5.2-7.2) | -12.8 to 11.8 | 100 (99–100) |
| 14 | -0.7 (-1.3 to -0.1) | 0.7 (0.1-1.3) | 4.1 (3.3-5.2) | -8.7 to 7.3 | 100 (99–100) |

In contrast, the standard deviation (SD) of the differences and the corresponding limits of agreement (LOA) widened substantially as data segment duration decreased (Table 4, Figure 3D). The median SD of differences was 12.5 sympathetic vasomotion score units at ten minutes, corresponding to LOA of -25.4 to 23.5 score units. At four minutes the SD of differences was 28.4 score units, with LOA of -56.0 to 54.6 score units, and at one minute the SD of differences was 63.2 score units with LOA of -119.3 to 130.3 score units. In the representative Bland-Altman plots (Figure 3A through 3C), differences were distributed around the zero-difference line across the range of sympathetic vasomotion scores without a clear pattern suggesting proportional bias, and both CDNx and SDNx kidneys and all ablation rounds contributed to the spread.

To place this scatter in context, individual agreement was also expressed against an absolute tolerance of 25 sympathetic vasomotion score units, a quarter of the 100-point separation between innervated and denervated kidneys at baseline (Table 4). The proportion of kidney-timepoints whose shortened data segment score fell within this tolerance of its own 15-minute score was 31 percent at one minute, 55 percent at three minutes, 65 percent at four minutes, 72 percent at five minutes, 84 percent at seven minutes, and 96 percent at ten minutes. Unlike group-level agreement, which changed little beyond four minutes, individual-level precision continued to improve throughout the range of durations examined.

### Decision Making and Practical Minimum Duration

Decision performance was assessed directly by asking whether shortened data segments supported the same conclusions as the full recordings (Table 5, Figure 4). At baseline, the CDNx kidney scored above its SDNx counterpart in 10 of 10 animals with the 15-minute recordings. Correct paired classification was 90 percent with one- and two-minute data segments and 95 to 100 percent from three minutes onward. The area under the receiver operating characteristic curve (AUC) for identifying the innervated kidney from a single score, without reference to the contralateral kidney, rose from 0.77 at one minute to 0.82 at four minutes and 0.84 at five to ten minutes, compared with 0.83 for the full 15-minute recordings.

**Fig. 4.**
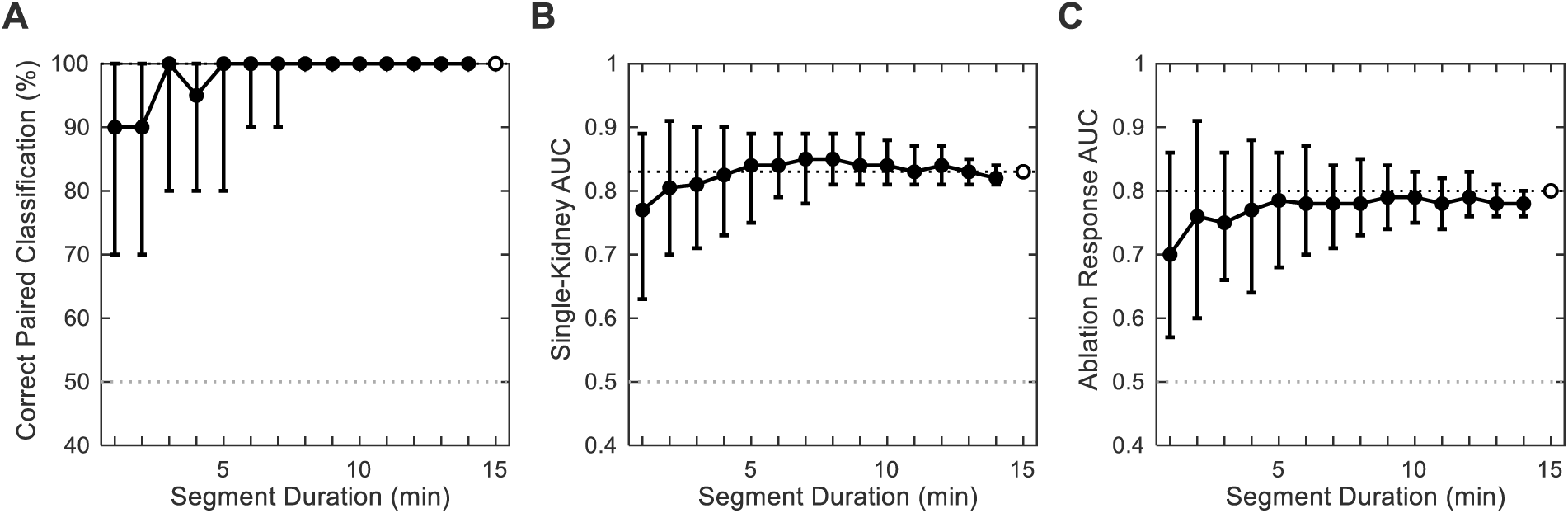
Decision performance of shortened data segments. (A) Correct paired classification: the percentage of animals in which the catheter-based denervation (CDNx) kidney, innervated at baseline, scored higher than the surgically denervated (SDNx) kidney of the same animal at baseline. (B) Single-kidney classification: the area under the receiver operating characteristic curve (AUC) for separating CDNx from SDNx kidneys by a single baseline score, without reference to the other kidney. (C) Ablation response: the AUC for separating the change in the CDNx kidney’s score, from baseline to after the fourth (final) ablation, from the change in the untreated SDNx kidney over the same interval. For data segment durations of 1 to 14 minutes, lines show the median across 100 sets of randomly placed data segments and error bars the 2.5th to 97.5th percentiles. Baseline and post-ablation data segments were drawn independently from their own recordings. Open markers at 15 minutes, extended as dotted horizontal lines, show performance with the full 15-minute recordings. Gray dotted lines mark chance. Correct paired classification was 90 percent with 1- and 2-minute data segments and 95 to 100 percent from 3 minutes onward (15 minutes: 100 percent). Single-kidney AUC was 0.82 at 4 minutes (15 minutes: 0.83), and ablation-response AUC 0.77 at 4 minutes (15 minutes: 0.80). n = 10 swine.

**Table 5:** Decision making metrics for shortened data segments as a function of recording length. Correct paired classification is the percentage of animals in which the catheter-based denervation (CDNx) kidney scored above the surgically denervated (SDNx) kidney at baseline. Single-kidney AUC is the area under the receiver operating characteristic curve for separating CDNx from SDNx baseline scores without reference to the contralateral kidney. Ablation-response AUC separates the change in the CDNx score, from baseline to after the second or the fourth ablation, from the change in the untreated SDNx kidney over the same interval. Values are the median over 100 randomized passes (2.5th to 97.5th percentile); at 15 minutes the full recording was used and a single value is given. Decision performance approached its 15-minute value by three to five minutes for all metrics.

| Duration (min) | Correct paired classification (%) | Single-kidney AUC | Ablation-response AUC, round 2 | Ablation-response AUC, round 4 |
| --- | --- | --- | --- | --- |
| 1 | 90 (70-100) | 0.77 (0.63-0.89) | 0.69 (0.54-0.83) | 0.70 (0.57-0.86) |
| 2 | 90 (70-100) | 0.80 (0.70-0.91) | 0.69 (0.58-0.84) | 0.76 (0.60-0.91) |
| 3 | 100 (80-100) | 0.81 (0.71-0.90) | 0.71 (0.62-0.82) | 0.75 (0.66-0.86) |
| 4 | 95 (80-100) | 0.82 (0.73-0.90) | 0.73 (0.64-0.81) | 0.77 (0.64-0.88) |
| 5 | 100 (80-100) | 0.84 (0.75-0.89) | 0.72 (0.62-0.80) | 0.78 (0.68-0.86) |
| 6 | 100 (90-100) | 0.84 (0.79-0.89) | 0.73 (0.67-0.79) | 0.78 (0.70-0.87) |
| 7 | 100 (90-100) | 0.85 (0.78-0.89) | 0.74 (0.68-0.78) | 0.78 (0.71-0.84) |
| 8 | 100 (100-100) | 0.85 (0.81-0.89) | 0.75 (0.70-0.79) | 0.78 (0.73-0.85) |
| 9 | 100 (100-100) | 0.84 (0.81-0.89) | 0.74 (0.71-0.79) | 0.79 (0.74-0.84) |
| 10 | 100 (100-100) | 0.84 (0.81-0.88) | 0.74 (0.71-0.78) | 0.79 (0.75-0.83) |
| 11 | 100 (100-100) | 0.83 (0.81-0.87) | 0.75 (0.71-0.78) | 0.78 (0.74-0.82) |
| 12 | 100 (100-100) | 0.84 (0.81-0.87) | 0.75 (0.73-0.77) | 0.79 (0.76-0.83) |
| 13 | 100 (100-100) | 0.83 (0.81-0.85) | 0.75 (0.72-0.77) | 0.78 (0.76-0.81) |
| 14 | 100 (100-100) | 0.82 (0.81-0.84) | 0.75 (0.73-0.76) | 0.78 (0.76-0.80) |
| 15 | 100 | 0.83 | 0.75 | 0.80 |

The ablation response, defined as the change in the CDNx score from baseline to the final ablation relative to the change in the untreated SDNx kidney over the same interval, was recovered less completely at every duration. The AUC was 0.70 at one minute, 0.77 at four minutes, and 0.79 at ten minutes, against 0.80 with the full recordings (Table 5, Figure 4C). Because this measure was limited at 15 minutes as well, its ceiling reflects variability of the untreated kidney between recordings rather than data segment duration alone.

Group-level agreement and decision performance therefore converge on approximately four minutes as the practical minimum data segment duration for comparisons between groups and for paired, within-individual decisions. At four minutes the relationship with the 15-minute sympathetic vasomotion score was preserved, with high correlation and ICC values and substantially lower error than durations below four minutes; the group-level trajectory across ablations was reproduced; and the decisions made from the full recordings were largely retained. Individual-level precision is a separate question, and by an absolute tolerance of 25 score units it continued to improve well beyond the knee, reaching 96 percent of kidney-timepoints only at approximately ten minutes.

## Discussion

This study evaluated the ability of shorter data recordings to reliably reflect sympathetic vasomotion scores derived from 15-minute reference data. Four-minute data segments preserved strong group-level agreement with reference 15-minute sympathetic vasomotion scores, maintained the statistical power needed to distinguish innervated from denervated renal sympathetic states, and reproduced the denervation decisions made from the full recordings. Thus, four-minute data segments may provide sufficient accuracy for cohort analysis and intraprocedural feedback for renal denervation.

Individual-level precision, however, continued to improve with recording length, and approximately ten-minute recordings were required before individual scores reliably fell within a quarter of the difference that separates an innervated from a denervated kidney. This higher degree of precision may be particularly important for stratifying patients based on resting renal sympathetic outflow preprocedurally.

The movement towards physiologically informed renal denervation echoes the coronary revascularization space, where decades of anatomic targeting of stenoses have given way to functional stenosis assessment with measures like fractional flow reserve and instantaneous wave-free ratio (Tonino et al., 2009; Min et al., 2012; Davies et al., 2017). Physiological recordings of four minutes are similar in length to those used for fractional flow reserve, but certainly longer than the few seconds of data needed to calculate the instantaneous wave-free ratio (Toth et al., 2016). That said, unlike coronary interventions, the target pathology for renal denervation is not visible angiographically, and this lack of anatomic feedback greatly increases the value of physiological validation for interventionists performing renal denervation.

The difference seen between group-level and individual-level reliability remains important. Group-level analyses primarily depend on central tendencies and relationships across conditions, and the random variations in short individual observations are offset by one another. This likely explains why four-minute recordings retained the statistical findings seen with the 15-minute recordings. From a group-level perspective, short sympathetic vasomotion measurements could be especially useful for future studies evaluating new renal denervation devices or comparing new ablation strategies with established devices. In these settings, the goal is often to determine whether a device or technique produces a measurable reduction in renal sympathetic activity across experimental groups, and the present data suggest that four-minute recordings preserve these group-level relationships.

Individual-level interpretation is more stringent because each shortened data segment must approximate its corresponding 15-minute data segment. Bland-Altman analysis showed that the main effect of shortening the data segment was not the introduction of large systematic bias, but rather an increase in the spread of individual differences. This suggests that shorter recordings do not consistently overestimate or underestimate sympathetic vasomotion, but instead become less representative of the full 15-minute data segment on an observation-by-observation basis.

At the individual level, the appropriate recording duration should be considered in the context of the clinical question and procedural environment. Intraprocedurally, each measurement can be interpreted against the same individual’s baseline or other kidney, so the relevant comparison is paired, and a four-minute recording appears sufficient. This is supported directly by the decision analysis, in which paired classification and the ablation-response comparison approach their 15-minute values by four minutes. Shortening data collection to approximately four minutes would improve feasibility, reduce procedural time and cost, and support real-time feedback in the cardiac catheterization laboratory.

Preprocedural measurements intended to stratify patients into responders and non-responders may require longer recordings. Unlike in the intraprocedural environment, these scores have no paired comparator, and the distribution of sympathetic vasomotion scores in patients with high or low renal sympathetic activity is not yet known. The relevant metric thus is the absolute precision of a single score, which reached 96 percent of observations within 25 score units only at approximately ten minutes. The time constraints in the preprocedural outpatient setting are much more relaxed than those in the procedural suite, and thus a ten-minute recording may be feasible as long as the technical challenges related to acquisition of ten minutes of continuous renal arterial blood flow velocity data can be overcome. This stable baseline could then be used to contextualize intraprocedural changes and support future studies evaluating whether sympathetic vasomotion helps identify patients most likely to respond to renal denervation.

The observation that systematic bias remained relatively low while variability increased with shorter data segment durations likely reflects underlying physiology. Muscle sympathetic nerve activity is dynamic and time-varying, but reproducible over longer recordings across many years (Fagius and Wallin, 1993; Notay et al., 2016). The temporal variability in sympathetic outflow may therefore produce outsize fluctuations in shortened data segments that are smoothed by longer recording segments. This may explain why four-minute recordings still preserved the coarser physiologic distinction between innervated and denervated states but ten-minute recordings were needed for more granular individual-level agreement.

Several limitations should be considered. First, the 15-minute data segment was treated as the reference, but it is not an absolute gold standard for sympathetic outflow, and future studies should link direct measures of sympathetic nerve activity to sympathetic vasomotion. In addition, because the shortened data segments were sampled from the 15-minute reference data, agreement between the shortened and 15-minute sympathetic vasomotion scores is higher than what would be observed with fully independent recordings. Additionally, these studies in anesthetized swine without hypertension certainly do not reflect the rich variability of pathophysiology in human hypertension. Finally, randomized segment selection estimates the reliability of shortened data recordings across many possible windows, but it does not determine whether specific portions of the recording are systematically more stable or more informative. Such analysis could inform adaptive approaches that assess the diagnostic quality of the signal and continue recording until a prespecified reliability criterion is met.

In summary, shortened data segments demonstrate strong group-level agreement with sympathetic vasomotion scores obtained from 15-minute data recordings. Paired comparisons and the decisions derived from them are also robust at short recording lengths while individual-level precision increases with recording duration. This analysis suggests four-minute recordings as a practical minimum duration for sympathetic vasomotion for group-level statistical inferences and for individualized intraprocedural feedback whereas ten-minute recordings may be the minimum duration for preprocedural assessment. As always, clinical judgement is paramount in balancing the efficiency gained by shortening data acquisition against the need for confidence in the measure itself.

**Perspectives: Sympathetic vasomotion, a novel marker of sympathetic outflow, can be evaluated at the group level and for paired intraprocedural decisions with data segments as short as four minutes, roughly a quarter of the recording duration used previously, while approximately ten minutes remains preferable when assessing preprocedural renal sympathetic tone at the individual level.**

## Generative AI Statement

The authors acknowledge the use of Claude Code for assistance with MATLAB code development and data visualization workflows. The authors also utilized ChatGPT 5.5 to enhance the clarity, flow, and grammatical correctness of the manuscript. All modifications were reviewed and approved by the authors.

